# Malnutrition disrupts adaptive immunity during visceral leishmaniasis by enhancing IL-10 production

**DOI:** 10.1101/2024.06.06.597776

**Authors:** Laís Amorim Sacramento, Claudia Lombana, Phillip Scott

## Abstract

Protein-energy malnutrition (PEM) is a risk factor for developing visceral leishmaniasis (VL). However, the impact on adaptive immunity during infection is unknown. To study the effect of malnutrition on chronic VL, we used a polynutrient-deficient diet (deficient protein, energy, zinc, and iron), which mimics moderate human malnutrition, followed by *Leishmania infantum* infection. The polynutrient-deficient diet leads to growth stunting and reduced mass of visceral organs. Malnourished-infected mice harbored more parasites in the spleen and liver, had a reduced number of T lymphocytes, reduced production of IFN-γ by T cells, and exhibited enhanced IL-10 production. To test whether IL-10 blockade would lessen disease in the malnourished mice, we treated infected mice with monoclonal antibody α-IL-10R. α-IL-10R treatment reduced the parasite number of malnourished mice, restored the number of T cells producing IFN-γ, and enhanced hepatic granuloma formation. Our results indicate that malnutrition increases VL susceptibility due to a defective IFN-γ-mediated immunity attributable to increased IL-10 production.

**Author Summary:** Malnutrition contributes to the development of VL. Despite the advances regarding this association, how malnutrition affects the adaptive immune mechanisms in VL is still unclear. We found that malnutrition disrupts the ability to control parasite replication in the spleen and liver in VL due to defective IFN-γ-mediated immunity, reduced hepatic granuloma formation, and enhanced IL-10 production. Blocking IL-10R signaling restored the protective mechanisms to control parasite replication in the malnourished mice without interfering with the undernutrition state. Thus, we demonstrate that malnutrition disrupts the adaptive immunity against VL due to an aberrant IL-10 production. Understanding the association between malnutrition and VL will provide insights into therapeutic approaches.

## Introduction

Protein-energy malnutrition (PEM) is the most frequent type of malnutrition, affecting at least 800 million people worldwide, especially children and elderly individuals [1], and is the most common cause of immunodeficiency worldwide [2]. Deficiency in macronutrients and/or micronutrients induces impaired immunity and consequently increases susceptibility to a variety of pathogens [3][4]. Visceral leishmaniasis (VL) is one of the infections whose risk is increased by malnutrition [5,6]. Murine models of moderate PEM demonstrate that malnutrition causes early dissemination of leishmania parasites to visceral organs due to defective innate immunity [7–9]. Despite substantial efforts to define the mechanisms between malnutrition and VL, the immunopathogenesis of this association is not completely understood. This study aimed to determine the impact of malnutrition on adaptive immunity during VL. We took advantage of a well-established model of polynutrient deficiency that mimics moderate human malnutrition [10] to fill this gap in knowledge. We found that malnutrition increases the susceptibility to VL due to defective IFN-γ-mediated immunity and hepatic granuloma formation as a consequence of an aberrant IL-10 response.

## Results

### Malnutrition disrupts anti-parasitic immunity during visceral leishmaniasis

We first characterize the impact of the polynutrient-deficient diet (PND diet) on the nutritional status of our model by monitoring body weight, tail lengths, and the size of the spleen and liver, which are the target organs of VL. Mice fed the PND diet gained significantly less weight over time than mice fed the control diet, and there is no statistical difference between naïve and infected malnourished mice (Fig 1A-B). Consistent with previous reports, the growth curve observed in PND diet-fed mice is considered a moderate level of malnutrition in mice [10] and mimics malnutrition in humans [11]. At the end of 6 weeks of feeding, mice on the PND diet had shorter tails (Fig1C). There was a significant reduction in the spleen (Fig 1D) and liver (Fig 1E) weight in the PND diet-fed mice. These data suggest that a PND diet led to growth stunting and reduced visceral organ size.

**Figure 1.**
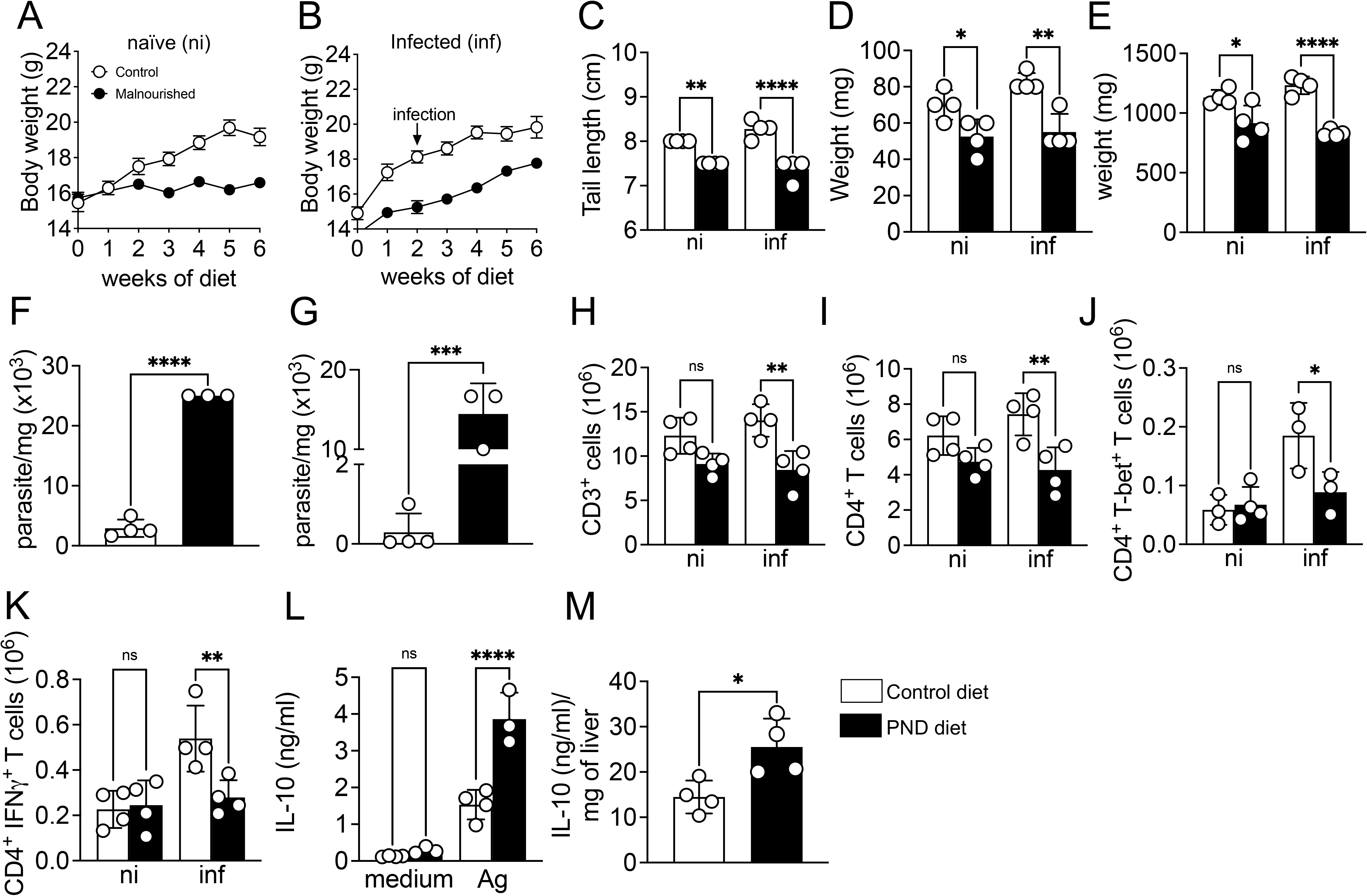
Malnutrition disrupts anti-parasitic immunity during visceral leishmaniasis. (A and B) Body weight of naive or *L. infantum*-infected control diet or polynutrient deficient diet (PND diet) over time. (C) Final tail lengths at 6 weeks of diet and 4 weeks post-infection (wpi). (D) Spleen and (F) liver weight. Parasite number in the (F) spleen and (G) liver of control diet and PND diet-fed mice at 4 weeks post-infection. Absolute number of (H) CD3^+^ cells, (I) CD4^+^ T cells, (J) CD4^+^ T cells expressing T-bet, and (K) CD4^+^ T cells expressing IFNγ in the spleen of control and PND diet-fed mice. Cells were gated based on their characteristic size (FSC) and granularity (SSC), singlet cells, live^+^, and CD45^+^ cells. (L) IL-10 levels in the supernatants from splenocytes restimulated with *L. infantum* crude antigen or medium for 72 h at 4 wpi. (M) IL-10 levels in the liver homogenate supernatant at 4 wpi. The data are expressed as mean ± SEM and represent two experiments (n = 4 mice/group). The statistical significance was calculated by one-way ANOVA or Student’s t-test (*p < 0.05, **p < 0.01, ***p < 0.001 and ****p < 0.0001).

To test whether malnutrition impacts the control of parasite replication, we performed limiting dilution for parasite quantification in the target organs of VL. Infected-malnourished mice harbored significantly more parasites at the spleen and liver at 4 weeks post-infection (wpi) (Fig 1F-G). We found a reduction in the number of CD3^+^ cells (Fig 1H) and CD4^+^ T cells (Fig 1I) in the spleen of infected-malnourished mice compared to infected-control. No significant differences were observed between non-infected malnourished versus control mice. The development of Th1-type immunity is critical for parasite control [12,13]. We found a reduction in the number of CD4^+^ T-bet^+^ T cells in the spleen of infected-malnourished mice compared with infected-control mice (Fig 1J). Consistently, the number of CD4^+^ T cells producing IFN-γ is significantly reduced in the spleen of infected-malnourished mice compared with infected-control mice (Fig 1K).

IL-10 is a key immunosuppressive factor in VL [14–18]. IL-10 levels were significantly higher in the supernatant of cultured cells from infected-malnourished mice versus the infected-control mice (Fig 1L). Consistently, liver samples from infected-malnourished mice exhibited increased amounts of IL-10 (Fig 1M). These data suggest that malnutrition results in a defect in the ability to develop Th1 cells following infection, which leads to an increased parasite burden, possibly due to a detrimental effect of the enhanced IL-10 production.

### IL-10R blockade restores anti-parasitic immunity

IL-10 is a pleiotropic cytokine that can influence VL development through multiple mechanisms [18–21]. To investigate the effect of IL-10 production in the infected malnourished mice, we treated mice with α-IL-10R mAb to block IL-10 signaling or with an α-IgG mAb as a control. Treatment was initiated at the time of infection and continued weekly. The treatment with α-IL-10R did not alter the body weight (Fig 2A-B) or tail length (Fig 2C) of malnourished mice. We observed a small increase in the spleen (Fig 2D) and liver (Fig 2E) weight of PND diet-fed mice treated with α-IL-10R compared with the control α-IgG mAb.

**Figure 2.**
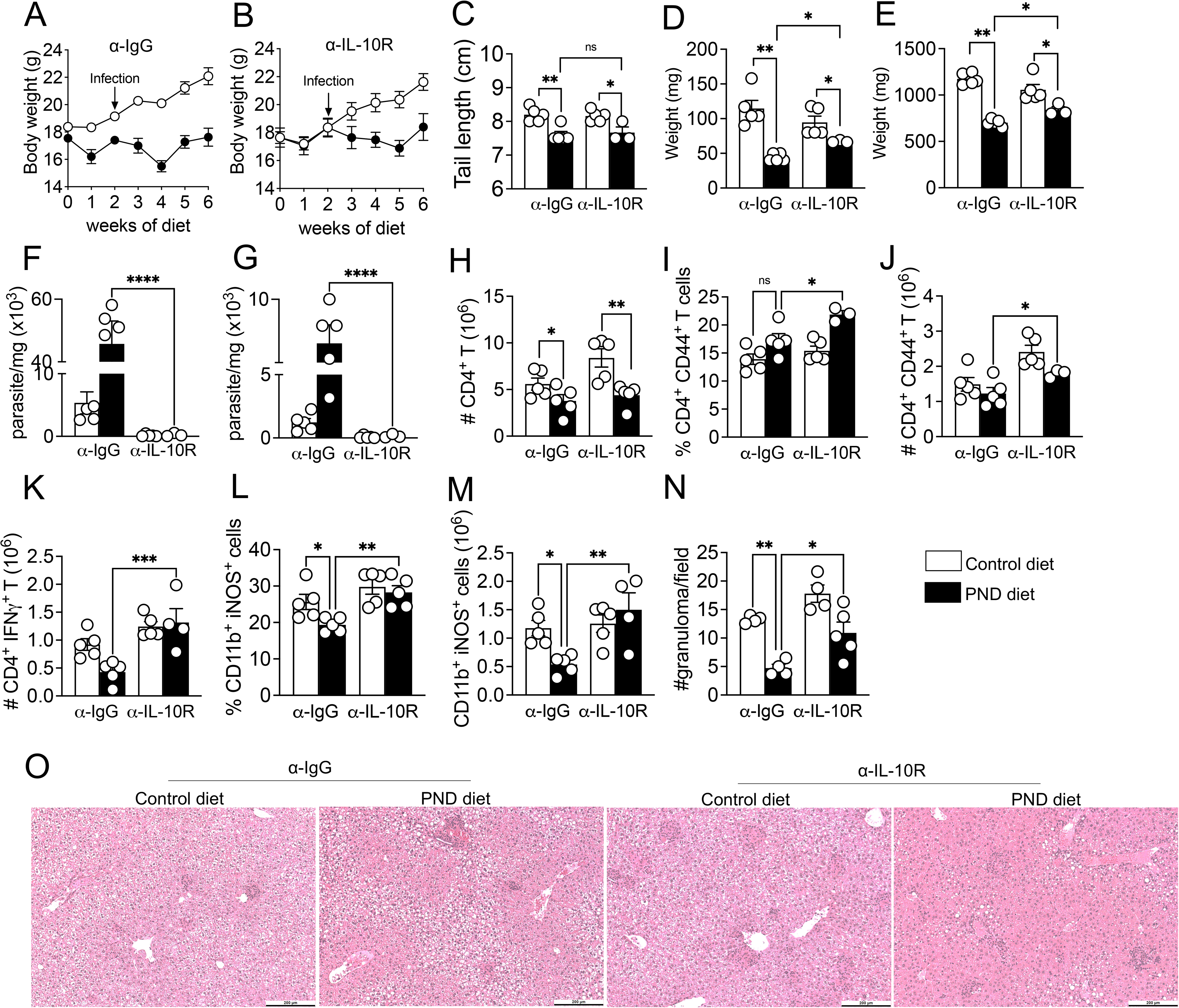
IL-10R blockade restores anti-parasitic immunity. *L. infantum*-infected mice fed with control diet or PND diet were treated with anti-IgG control or anti-IL-10R antibody and euthanized at 4 wpi. Treatment was initiated at the time of infection and continued weekly until the termination of the experiment. (A and B) Body weight of infected mice treated with (A) α-IgG control or (B) α-IL-10R antibody. (C) Tail lengths at 6 weeks of diet and 4 weeks post-infection (wpi). (D) Spleen and (E) liver weight. Parasite number in the (F) spleen and (G) liver. Absolute number of (H) CD4^+^ T cells, (I) percentage and (J) number of CD4^+^ T cells expressing CD44, (K) number of CD4^+^ T cells expressing IFNγ at the spleen of control and PND diet-fed mice treated with anti-IgG control or anti-IL-10R antibody. (L-M) Percentage and number of CD11b^+^ cells expressing iNOS at the spleen of infected control and PND diet-fed mice treated with α-IgG control or α-IL-10R antibody. Cells were gated based on their characteristic size (FSC) and granularity (SSC), singlet cells, live^+^, and CD45^+^ cells. T cells were gated on CD3^+^, CD4^+^, and myeloid cells on CD11b^+^, CD11b^+^ iNOS^+^ cells. (N-O) Representative pictures of H&E-stained liver sections (10x magnification) and bar graph of the number of granulomas per field. The data are expressed as mean ± SEM and represent two experiments (n = 3-5 mice/group). The statistical significance was calculated by one-way ANOVA (*p < 0.05, **p < 0.01, ***p < 0.001 and ****p < 0.0001).

As previously reported, α-IL-10R treatment reduced the parasite number in the spleen and liver of well-nourished mice [19]. Importantly, IL-10 signaling blockade reduced the parasite burden in the target organs of infected-malnourished mice (Fig 2F-G). We did not observe a difference in CD4^+^ T cell numbers in the spleen from α-IL-10R mAb treated mice compared with α-IgG control mice (Fig 2H). However, we found that α-IL-10R treatment increased the percentage and number of CD4^+^ CD44^+^ T cells (Fig 2I-J) and the number of CD4^+^ IFNγ^+^ T cells in the spleen of the infected-malnourished mice (Fig 2K).

iNOS is essential for nitric oxide production and control of parasites within macrophages [18–20]. We next sought to evaluate whether the IL-10R blockade interferes with the expression of iNOS in malnourished infected mice. The treatment with α-IL-10R mAb enhanced the percentage and number of CD11b^+^ cells expressing iNOS from the spleen of infected malnourished mice (Fig 2L-M).

The control of liver infection during experimental VL depends on granuloma formation [20]. We evaluated the liver sections to determine whether the enhanced IL-10 production in the infected-malnourished mice interfered with the hepatic granuloma generation. α-IL-10R mAb treatment increased the granuloma formation in the control mice, as previously reported [17]. Infected-malnourished mice had a reduced number of hepatic granulomas versus control mice. Importantly, the blockade of IL-10 signaling partially restored the number of granulomas in the liver of infected malnourished mice (Fig 2O). Our results indicate that malnutrition increases VL susceptibility due to defective activation and expansion of CD4^+^ IFNγ^+^ T cells and reduced hepatic granuloma formation, which is attributable to increased IL-10 production.

## Discussion

We found that malnutrition increases VL susceptibility due to elevated IL-10 production that limits IFNγ-mediated immunity and hepatic granuloma formation. Malnutrition disrupts innate immune mechanisms and causes early visceralization of parasites during VL [7,9,10], affects the development of T cells in the thymus of young mice, and leads to alterations in the intestine [21]. Additionally, malnutrition affects T cell-mediated responses and granuloma formation [22–24], and we found that α-IL-10R treatment restored activation and expansion of T cells producing IFNγ in the malnourished-infected mice without altering the undernutrition observed in the mice. The restoration of iNOS expression with the α-IL-10R treatment can be a consequence of enhanced CD4^+^ IFNγ^+^ T cells or a direct effect on the myeloid cells [15,25]. In human VL, increased IL-10 is associated with VL progression [15].

In agreement with our data, high levels of systemic IL-10 have been reported in malnourished mice challenged intraperitoneally with bacterial LPS [26–28]. Malnourished individuals [29,30] and murine models of malnutrition have increased circulating levels of inflammatory cytokines [31], which is attributed to a dysfunctional intestinal barrier and bacterial translocation [31]. Based on this, we hypothesize that the increased IL-10 production observed in the malnourished model might result from an excessive systemic inflammatory process. As a side effect, IL-10 suppresses immune mechanisms that control parasites. This idea is supported by the fact that high levels of IL-10 in sepsis are critical in controlling inflammatory response [32,33]. In support, VL patients have a systemic overproduction of inflammatory mediators [34,35], and the high levels of IL-10 limit tissue damage despite generating a permissive environment for parasite replication [36–38][14]. In summary, our findings indicate that malnutrition disrupts Th1 immunity and hepatic granuloma during VL due to enhanced IL-10 production.

## Material and Methods

### Mice and diets

4-week-old female C57BL/6 mice were purchased from Charles River. Mice were grouped in cages of 4-5 and were maintained in a specific pathogen–free facility with free access to water, nesting material, and housed at a temperature of 21°C at the University of Pennsylvania Animal Care Facilities. Mice received a control standard chow (17% protein, 100 ppm Iron, 30 ppm zinc - Teklad diet, #99103) or an isocaloric but protein, iron, and zinc-deficient diet (3% protein, 10 ppm Iron, 1 ppm zinc - Teklad diet, #99075). After 2 weeks of diet, mice were divided into two subgroups, one of which was infected with *L. infantum*. Polynutrient deficient diet-fed (PND diet) mice received 90% of the consumed food of the control mice over weeks calculated based on the food ingestion (g/mouse/day) of the control diet group obtained every 72 hours. The body weight of mice was monitored once a week. All animals were used in accordance with the recommendations in the Guide for the Care and Use of Laboratory Animals of the National Institutes of Health and the guidelines of the University of Pennsylvania Institutional Animal Use and Care Committee. The protocol was approved by the Institutional Animal Care and Use Committee, University of Pennsylvania Animal Welfare Assurance.

### Parasite culture and infection

*L. infantum* (LLM-320) was cultured in Schneider medium with 20% heat-inactivated fetal bovine serum, 5% penicillin, and streptomycin. The mice were intravenously infected via the retro-orbital plexus with 10^7^ *L. infantum* metacyclic enriched promastigotes in 100 μL of PBS. A quantitative limiting dilution assay determined parasite burdens.

### Splenocyte culture and flow cytometry

Spleen cells were stimulated with or without *L. infantum* crude antigen for 72h of culture at 37^°^C in 5% CO_2_, and IL-10 levels were accessed by ELISA. For intracellular staining, single-cell suspensions were incubated with PMA (50 ng/mL), ionomycin (500 ng/mL), and Brefeldin A (10 μg/mL) for 4h. Cells were stained with LIVE/DEAD Fixable Aqua Dead Cell Stain Kit and subsequently incubated with anti-CD16/CD32 and 10% rat-IgG1. For surface staining, cells were incubated with monoclonal anti-CD45, anti-CD3, anti-CD4, anti-CD11b, and anti-iNOS. For intracellular staining, cells were permeabilized with a Cytofix/Cytoperm kit according to the manufacturer’s guide and stained with anti-T-bet and α-IFNγ. Data were collected using LSRIII Fortessa and analyzed using FlowJo.

### In vivo α-IL-10R antibody treatment

Mice were treated intraperitoneally twice a week with 500 μg of α-mouse IL-10R antibody or rat anti-IgG1 isotype control. Treatment was initiated at the time of infection and continued weekly until the termination of the experiment at 4 weeks post-infection.

### Statistical analysis

Data are shown as means ± SEM. Statistical significance was determined using the two-tailed unpaired Student’s t-test or one-way ANOVA. All statistical analysis was calculated using GraphPad Prism Software. Differences were considered significant when *p < 0.05, **p ≤ 0.01, ***p ≤ 0.001, ****p < .0001.

## Acknowledgments

We thank the Comparative Pathology Core (CPC). This work was supported by National Institutes of Health grant RO1-AI150606 (PS). The funder had no role in study design, data collection and analysis, decision to publish, or preparation of the manuscript.

## Author contributions

Conceptualization: PS, LAS; Methodology: LAS, CGL; Validation: PS, LAS, CGL; Formal Analysis: LAS; Investigation: PS, LAS; Resources: PS; Writing - Original Draft: LAS; Writing - Review & Editing: PS, LAS; Visualization: LAS; Project administration: PS; Experimental work: LAS, CGL; Funding Acquisition: PS; and Supervision: PS.

